# Encapsulated mononuclear stem cells: paracrine action for the treatment of acute myocardial infarction

**DOI:** 10.1101/448415

**Authors:** Santiago A. Tobar, M. Andrades, V Olsen, D. Silvello, A Phaelante, A Lopes, A Biolo, U Matte, LE Rohde, NO Clausell

**Affiliations:** Laboratory of Cardiovascular Research, Experimental Research Center Hospital de Clínicas de Porto Alegre, Rua Ramiro Barcelos, 2350, Porto Alegre, RS 90035-003 Brazil; Post Graduation Program in Medical Science: Cardiology and Cardiovascular Science, Universidade Federal do Rio Grande do Sul, Brazil; Gene Therapy Center, Experimental Research Center, Hospital de Clínicas de Porto Alegre, Brazil

**Keywords:** Myocardial Infarction., Bone-Marrow Mononuclear Cells., Stem Cells., Cardiac Regeneration., Cell Therapy., Sodium Alginate., Paracrine action.

## Abstract

Cell therapy is considered as a treatment option for acute myocardial infarction (AMI). Released molecules by cells paracrine action may promote tissue regeneration. Therefore we used bone-marrow mononuclear cells (BMMNCs) from GFP+ Wistar rats encapsulated in sodium alginate for AMI treatment. Animals were randomly allocated into groups – empty (EC); BMMNC capsules; or sham. AMI was induced by occlusion of left anterior artery and capsules were delivered intrathoracically. Troponin I was measured 24h after AMI and echocardiography was performed at 48h and 7d after AMI. On day 7 animals were euthanized and their hearts were harvested. Tissue levels of TNF-α, IL-6, IL-10, cleaved caspase-3, and catalase were measured. Technical procedures were performed by blinded operators. There was no difference in either heart morphofunctional parameters or biochemical analysis between AMI groups. We conclude that the paracrine effects of BMMNCs lacks efficacy to modulate events associated with AMI in the rat.

## Introduction

Coronary artery disease (CAD) [1] is responsible for approximately 17.5 million deaths a year worldwide [2,3]. Acute myocardial infarction (AMI) is the main endpoint of CAD as a consequence of coronary artery occlusion. The decreased supply of nutrients and oxygen to cardiomyocytes leads to inflammation, oxidative stress, cell death, and impairment of cardiac function. The post-MI healing process includes cardiac remodeling, which is characterized by fibrotic tissue formation, hypertrophy, and increased ventricular chamber volume; progression of remodeling is known to contribute to heart failure [4,5]. Regardless of treatment improvements in the last 30 years, mortality is still significant, and the only curative option is heart transplantation [4]. Consequently, artificial organs and regenerative interventions have emerged as promising approaches in experimental studies [4,5].

Stem cell-based therapy for cardiovascular diseases has been widely studied, due to the ability of these cells to transdifferentiate into cardiomyocytes [6,7], fuse with resident cells [8-10], and release factors in a paracrine mechanism that may activate cardiac progenitor cells or control inflammation [11,12]. However, inconsistency in findings across clinical trials has emerged as the main barrier to establishment of therapy [13,14]. Misapplication of stem cells could be the main reason behind the discrepancies observed. For instance, as reviewed by Segers and Lee (2008), many studies have delivered stem cells by intracoronary route, which allows cells to be washed by blood flow so the few cells actually remaining in the heart have to deal with a hostile, inflammatory milieu, making regeneration very difficult [5,15].

Encapsulation of cells is an interesting method to study their paracrine action, as it prevents stem cells from fusing with resident cells, but still allows hormonal communication and flow of nutrients and metabolites. Additionally, encapsulated stem cells are protected from the host immune response [16,17].

The goal of this study was to evaluate the paracrine effect of encapsulated bone marrow-derived mononuclear cells (BMMNCs) on cardiac remodeling and inflammatory response in a murine model of acute myocardial infarction.

## Methods

### Animals

Four (04) male GFP^+^ Wistar rats and 67 male GFP^−^ Wistar rats (weight 337±31 g, 2 months) were used in this study. The protocol was approved by the Hospital de Clínicas de Porto Alegre Animal Care and Use Committee and registered with number 10-0246. All animals had access to standard rat chow and water ad libitum and were kept under 12:12-h light/dark cycles and controlled temperature and humidity. All procedures were performed to ensure minimal suffering, with anesthesia and postoperative analgesia. Animals were euthanized by exsanguination under deep anesthesia with ketamine (100 mg/kg) and xylazine (10 mg/kg).

### Groups

Animals were randomly allocated across study groups. AMI was induced in 61 animals divided into two treatment groups: those receiving encapsulated BMMNCs (n=31) and those receiving empty capsules (EC) (n=30). In addition, a sham group (n=6), which underwent surgery without coronary ligation and received saline as treatment, was included in the study as control.

### Acute Myocardial Infarction

AMI was induced as described by Pfeffer et al. (1979) [18]. Briefly, animals were weighed, anesthetized with ketamine (100 mg/kg i.p.) and xylazine (10 mg/kg i.p.), maintained on 37 °C hot plate, and artificially ventilated using a Harvard Apparatus small-animal ventilator pumping 2 mL of air at a rate of 88 breaths per minute (bpm) during surgery. A small incision was made in the thoracic region and the pectoralis muscles and ribs were retracted to expose the heart. The pericardium was broken and then the left descending coronary artery was permanently occluded by suturing with 6-0 Mononylon. After coronary ligation, the chest was closed and analgesia was provided by intramuscular injection of bupivacaine (1 mg/kg). The average operative time was 14 min. After surgery, all animals were kept in a controlled-temperature and oxygen-enriched environment until recovery.

### Echocardiography

Echocardiography was performed in all animals 48 h and 7 d after AMI. Before echocardiography, animals were anesthetized with ketamine (100 mg/kg i.p.) and xylazine (10 mg/kg i.p.) and the chest area was shaved. Scans were performed by a blinded technician using an EnVisor ultrasound system (Philips, Andover, MA, USA) with a 12-3 MHz transducer and 2 cm depth. Left ventricle diastolic diameter (LVDD) and left ventricle systolic diameter (LVSD) were measured in M-mode. Infarct size was measured by the akinetic/hypokinetic arc (AHA)/total endocardial perimeter ratio in three cross-sectional views (basal, at the border of the cusps; medial, at the level of the papillary muscles; and in the apical region) according to the equation %AMI= (AHA/EPt)x100, as previously described [19].

### BMMNC encapsulation

Femoral bone marrow from four non-operated GFP^+^ Wistar rats was flushed into DMEM medium and centrifuged at 800×*g* for 10 min. Pelleted cells were resuspended in 3 mL PBS, mixed to 3 mL Ficoll, and centrifuged at 800×*g* for 20 min. Mononuclear cells layer were collected, washed three times in PBS, and stored in DMEM medium. Cells were counted using Trypan blue to check for viability at time 0 and 24h after withdrawn [20,21].

BMMNCs were mixed with 1.5% sodium alginate (NOVAMATRIX, Norway) (5.0×10^6^ cells/mL) and 3 mL of solution were extruded in an air encapsulation unit (Nisco, Switzerland), assisted by a syringe pump, with flow adjusted to 20 mL/h and the air jet adjusted to 10 L/min. The extruded drops fell into 125 mM CaCl_2_ to induce alginate polymerization. Encapsulated cells were washed in 0.1% of Poly-L-Lysine, then washed again in 1% manitol and a final wash in 0.1% of Sodium Alginate. The finished Alginate-Poly-L-Lysine (Algina-PLL) capsules was washed again in 1% manitol and kept in DMEM until use. The protein cut-off of Alginate-PLL is up to 80 kDa [22,23]. Before administration, all capsules (561.15±178.3 µm) were washed three times in PBS and 300 µL of capsules were applied into the thoracic cavity shortly after AMI induction [24-26].

### BMMNC viability

At day 7 after AMI, capsules containing GFP^+^ mononuclear cells were collected from animals. Briefly, animals were anesthetized with 10% isoflurane, exsanguinated, and the thoracic cavity was filled with 1× PBS through a small incision to resuspend the capsules, which were then withdrawn and transferred to a tube. Capsules were washed three times in 1× PBS and alginate beads were dissolved by 10 mM sodium citrate solution for 10 min. Supernatant was collected and centrifuged for 10 min at 800 rpm (FANEM Excelsa II 206 BL, São Paulo, Brazil) at room temperature. Pelleted cells were resuspended in 50 µL of 1× PBS and analyzed under a fluorescence microscope.

### Tissue preparation

Hearts were harvested immediately after euthanasia and cut into three sections: infarct area (decellularized fibrotic tissue), peri-infarct area (myocardial tissue in the border of the infarct area), and remote area (unaffected myocardial tissue). Sections were homogenized in PBS buffer (pH 7.2) containing 1% Triton X-100 and a protease-inhibitor cocktail (SigmaFast, 1 tablet/40 mL) at a 1:10 ratio (1 g tissue per 10 mL of homogenizing buffer). Homogenized tissues were then centrifuged at 12,000×*g* for 10 min at 4°C, and the protein content of the supernatant was determined [27].

### Interleukin analysis

Cardiac cytokines were measured using a customized multiplex Luminex kit (catalog #LCP0004M, lot #1639562A), in accordance with manufacturer recommendations. Briefly, 50 µL of homogenized tissue was mixed with activated beads containing interleukin (IL)-6, IL-10, or tumor necrosis factor alpha (TNF-α) antibodies and incubated for 16 h at 4°C, 550 rpm, in a rotatory shaker. Then, beads were washed three times and incubated with biotinylated antibody for 2 h at room temperature in a rotatory shaker at 550 rpm. After incubation, beads were washed three times and incubated with streptavidin R-phycoerythrin (RPE)-conjugated antibody. Cytokine concentrations were recorded in a LUMINEX χmax-200 system.

### Antioxidant enzyme

Catalase (CAT) activity was measured as previously described [28]. Briefly, the reaction was performed in 50 mM of sodium phosphate buffer (pH 7.0), 10 mM hydrogen peroxide, and 10 µL of total heart homogenate. Hydrogen peroxide consumption was monitored spectrophotometrically at 240 nm, and enzyme activity was calculated using the H_2_O_2_ molar coefficient (43.6 M^−1^ cm^−1^).

### Apoptosis

Cleaved caspase-3 was assessed by Western blotting as follows: 30 µg of homogenized tissue was diluted in Laemmli buffer (250 mM Tris, 8% SDS, 40% glycerol, 20% β-mercaptoethanol, 0.008% Bromophenol Blue, pH 6.8) to a final protein concentration of 1 µg/µL. The mixture was heated for 10 min at 70°C and then loaded into SDS-PAGE 12%. Electrophoresis was performed for 90 min at 120 V before transferring to polyvinylidene fluoride (PVDF) membranes (Millipore Corporation, MA) in transfer buffer (25 mM Tris, 190 mM glycine, 20% methanol, pH 8.3) in a semidry system (Bio-Rad Laboratories, CA). The membranes were blocked with 5% nonfat milk in TTBS buffer (TBS 1x, 0.1% Tween-20) for 1 h and incubated overnight with anti-cleaved caspase-3 (Asp175) antibody (1:100) (Cell Signaling – #9661). Glyceraldehyde-3-phosphate dehydrogenase (GAPDH) was used as the housekeeping protein for cleaved caspase-3 band normalization. GAPDH was assessed with anti-GAPDH antibody (Cell Signaling - #5174) (1:6,000).

After primary antibody incubation, the membrane was washed and then incubated with secondary antibody (anti-rabbit IgG conjugated to horseradish peroxidase, Millipore - #AP307P) for 2 h at room temperature for cleaved CASP-3 (1:3,000) and GAPDH (1:6,000).

Image acquisition was performed using an L-Pix Chime Molecular Imaging system (Loccus Biotecnologia, Brazil), and band densitometry was determined on ImageJ software environment. Results were expressed as arbitrary units (density of cleaved CASP-3/density of GAPDH).

### Statistical Analysis

All data was tested for normality using Shapiro-Wilk test and expressed as mean±SD. Comparisons between groups were made using an unpaired Student’s t-test for endpoint comparison between BMMNC treatment against empty capsules or two-ways ANOVA with Tukey’s post-hoc test for follow-up comparison between treatment groups (BMMNC and EC) and SHAM. For cause of death distribution analysis between treatment groups a Chi-square test was performed.

All analysis and graphics were done using GraphPad Prism software version 6.0.

## Results

A total of 67 animals were randomly assigned to the sham, EC, or BMMNC groups. The mortality rate was nearly 33% (22 animals). Eight animals died during surgery (≈40% of total deaths), 12 died up to 48 h after surgery (≈60% of total deaths), and 45 survived. Nevertheless, as the outcome of AMI surgery is highly variable, only those animals with an infarct area > 30% were included in the study, leading to the exclusion of 12 animals. Hence, only 33 animals were included, with a balanced distribution into experimental groups (EC, n=13; BMMNC, n=15, SHAM, n=5) (Supplementary Fig 1). It is also important to note that no difference in mortality rate was observed between groups, neither during surgery nor during recovery (48 h after AMI) (Supplementary Fig 2). Animals included in the study had the same infarct size (EC = 38.9%±6.4%; BMMNC = 43%±7.2%; p>0.18) and troponin-I levels (EC = 31±13.5 ng/mL; BMMNC = 36.6±10.6 ng/mL; p>0.23) in both groups 48 h after AMI (Fig 1). This is a highly relevant finding, because it highlights that similar infarct sizes were induced in both groups.

**Fig 1.**
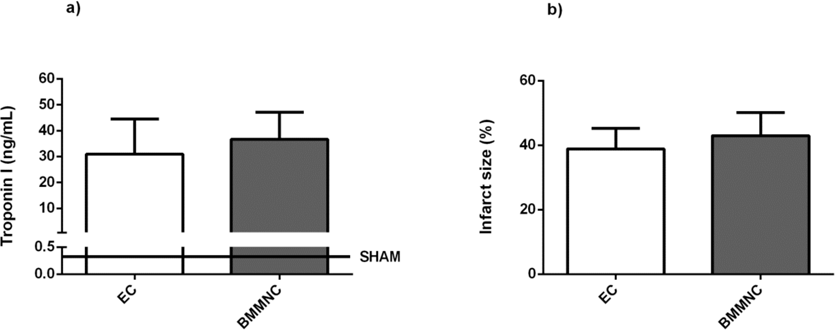
Assessment of heart damage. A) Heart damage was assessed 48 h after AMI by plasma troponin I measurement. B) Infarct size measured by echocardiography in animals receiving empty capsules (EC) or encapsulated bone marrow-derived mononuclear stem cells (BMMNCs). Data expressed as mean ± SD. Statistical significance accepted at p<0.05.

We assessed BMMNC viability after cell isolation, just before capsules implementation and on day 7 after AMI induction to ensure that any lack of effect would not be due to loss of cell viability. Cell viability at the time of BMMNC havesting was 82.7% and during the next 24h a viability loss rate was 11.8±6.7%. Therefore, at the time of capsules implementation at least 74% of BMMNC were still viable (Supplementary Fig 3). GFP^+^ BMMNCs withdrawn from depolymerized capsules were visualized in a fluorescence microscope. These cells were still viable at the end of the protocol (Fig 2).

**Fig 2.**
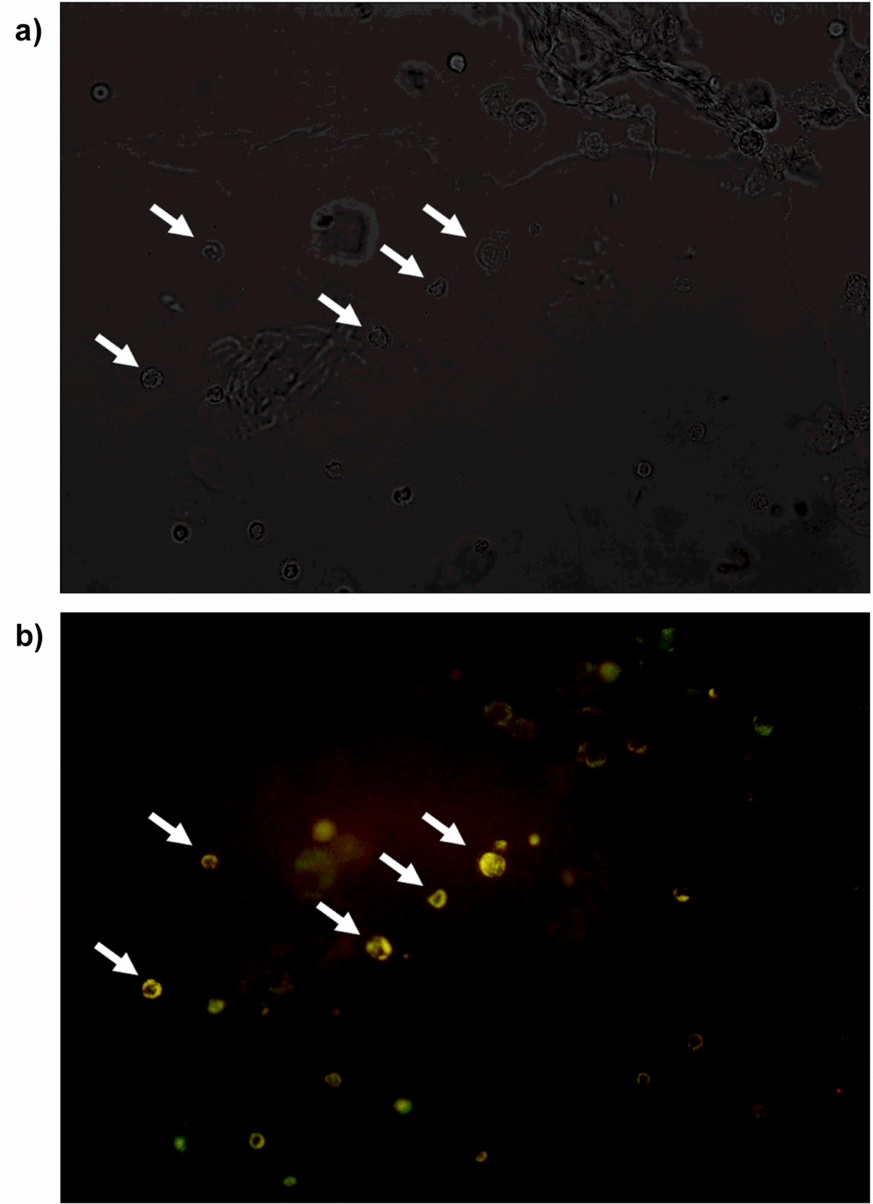
Cell viability 7 days after AMI. GFP+ cells collected from alginate capsules withdrawn from animals 7 days after AMI induction. Panels A and B represent GFP+ cells (arrows) in the same field, analyzed under visible light (A) and fluorescence microscopy (B) at 40x magnification.

Cardiac remodeling 7 days after AMI was the main outcome measure of treatment efficacy. As expected, both left ventricular end-diastolic and end-systolic diameter increased 7 days after AMI (p<0.01); however, BMMNC therapy did not change either parameter (p=0.56 and 0.34, respectively). Accordingly, anterior wall thickness in both systole and diastole decreased after 7 days of AMI (p=0.02 and 0.01, respectively). Again, the paracrine effect of BMMNCs was unable to prevent infarct wall remodeling (p=0.15 for systole and p=0.12 for diastole). The posterior wall remained unchanged (Table 1).

**Table 1.**
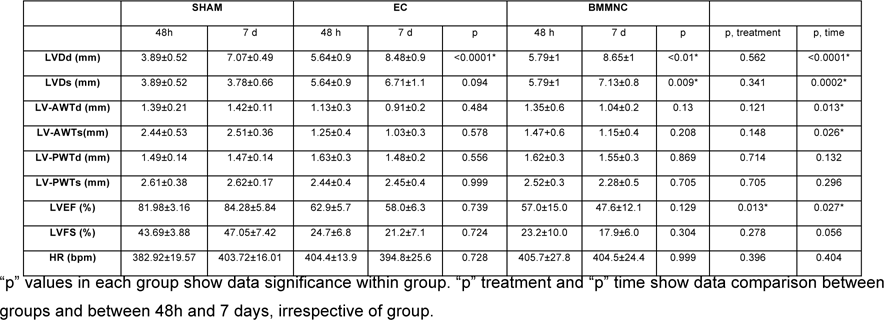
Echocardiography: LVDd, left ventricular diastolic diameter; LVDs, left ventricular systolic diameter; LVAWTd, left ventricular diastolic anterior wall thickness; LVAWTs, left ventricular systolic anterior wall thickness; LVPWTd, left ventricular diastolic posterior wall thickness; LVPWTs, left ventricular systolic posterior wall thickness; LVEF, left ventricular ejection fraction; LVFS, left ventricular fractional shortening; HR, heart rate.

|  | SHAM |  | EC |  |  | BMMNC |  |  |  |  |
| --- | --- | --- | --- | --- | --- | --- | --- | --- | --- | --- |
|  | 48h | 7 d | 48 h | 7 d | p | 48 h | 7 d | p | p, treatment | p, time |
| LVDd (mm) | 3.89±0.52 | 7.07±0.49 | 5.64±0.9 | 8.48±0.9 | <0.0001* | 5.79±1 | 8.65±1 | <0.01* | 0.562 | <0.0001* |
| LVDs (mm) | 3.89±0.52 | 3.78±0.66 | 5.64±0.9 | 6.71±1.1 | 0.094 | 5.79±1 | 7.13±0.8 | 0.009* | 0.341 | 0.0002* |
| LV-AWTd (mm) | 1.39±0.21 | 1.42±0.11 | 1.13±0.3 | 0.91±0.2 | 0.484 | 1.35±0.6 | 1.04±0.2 | 0.13 | 0.121 | 0.013* |
| LV-AWTs (mm) | 2.44±0.53 | 2.51±0.36 | 1.25±0.4 | 1.03±0.3 | 0.578 | 1.47±0.6 | 1.15±0.4 | 0.208 | 0.148 | 0.026* |
| LV-PWTd (mm) | 1.49±0.14 | 1.47±0.14 | 1.63±0.3 | 1.48±0.2 | 0.556 | 1.62±0.3 | 1.55±0.3 | 0.869 | 0.714 | 0.132 |
| LV-PWTs (mm) | 2.61±0.38 | 2.62±0.17 | 2.44±0.4 | 2.45±0.4 | 0.999 | 2.52±0.3 | 2.28±0.5 | 0.705 | 0.705 | 0.296 |
| LVEF (%) | 81.98±3.16 | 84.28±5.84 | 62.9±5.7 | 58.0±6.3 | 0.739 | 57.0±15.0 | 47.6±12.1 | 0.129 | 0.013* | 0.027* |
| LVFS (%) | 43.69±3.88 | 47.05±7.42 | 24.7±6.8 | 21.2±7.1 | 0.724 | 23.2±10.0 | 17.9±6.0 | 0.304 | 0.278 | 0.056 |
| HR (bpm) | 382.92±19.57 | 403.72±16.01 | 404.4±13.9 | 394.8±25.6 | 0.728 | 405.7±27.8 | 404.5±24.4 | 0.999 | 0.396 | 0.404 |

"p" values in each group show data significance within group. "p" treatment and "p" time show data comparison between groups and between 48h and 7 days, irrespective of group.

Observing LVEF alone revealed a significant decrease in the BMMNC-treated group as compared to the EC group (p=0.013), indicating that mononuclear cells could lead to deterioration of heart function. However, when the change in LVEF between 48 h and 7 d after AMI (LVEF_48h_ – LVEF_7d_) was analyzed, this difference between treatment groups did not remain significant (Fig 3)

**Fig 3.**
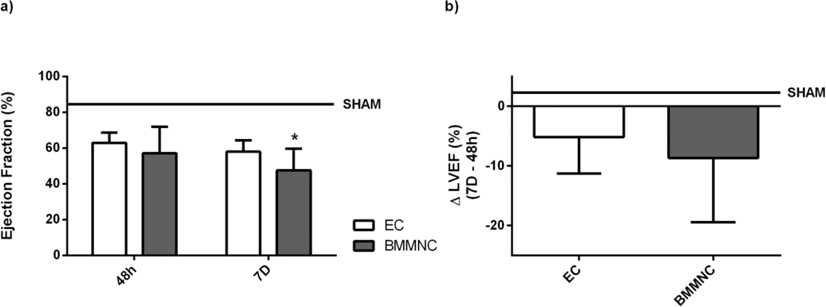
Ejection fraction. (A) Left ventricular ejection fraction of animals 48 h and 7 d after AMI induction. (B) Progression of ejection fraction from 48 h to 7 d post-AMI. Data expressed as mean ± SD. Statistical significance accepted at p<0.05.

We have also evaluated inflammatory response, catalase activity, and apoptosis as secondary outcome measures of treatment efficacy. Again, as expected, levels of the inflammatory cytokines TNF-α and IL-6 were increased in the infarct area of the left ventricle when compared to the peripheral area and remote (septum) areas (TNF and IL-6: p<0.01), but treatment was not associated with difference in any of these areas (TNF: p=0.216; IL-6: p=0.851); despite the apparent increase in TNF-α levels in the infarct area in the BMMNC group, this increase was not statistically significant (p=0.296). On the other hand, levels of the anti-inflammatory cytokine IL-10 remained the same in all areas for both groups (EC and BMMNC, Fig 4). In addition, we analyzed the activity of catalase to evaluate the ability to overcome the oxidative stress that affects inflamed tissues. Again, treatment with encapsulated mononuclear stem cells was unable to improve heart antioxidant capacity (Fig 5).

**Fig 4.**
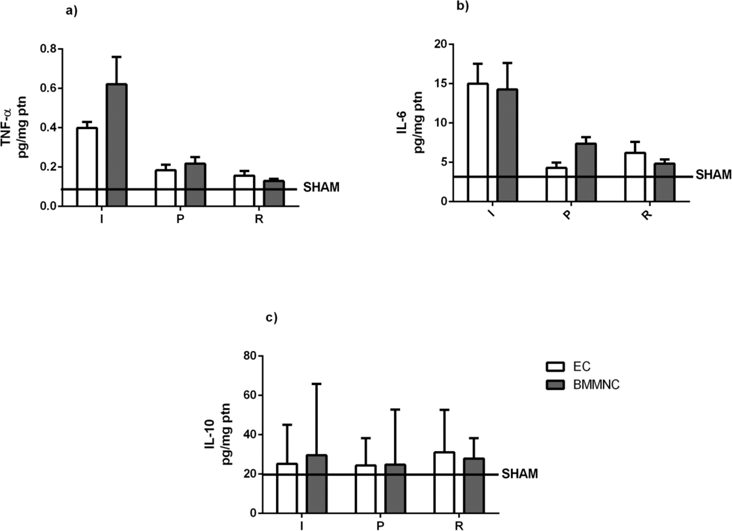
Cytokine levels. Tissue levels of TNF-α (A), IL-6 (B), and IL-10 (C) in three different heart regions. I, infarct region; P, peri-infarct region; R, remote (septum) region. White bars represent animals treated with empty capsules, while gray bars represent animals treated with encapsulated BMMNCs. Data expressed as mean ± SD. Statistical significance accepted at p<0.05

**Fig 5.**
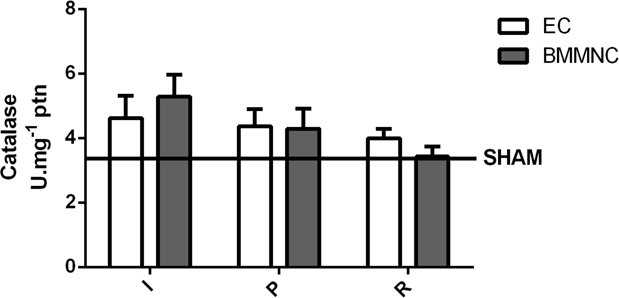
Catalase activity. Catalase activity in three different heart regions. I, infarct region; P, peri-infarct region; R, remote (septum) region. White bars represent animals treated with empty capsules, while gray bars represent animals treated with encapsulated BMMNCs. Data expressed as mean ± SD. Statistical significance accepted at p<0.05

Finally, we assessed the active form of caspase-3 (cleaved CASP-3) by Western blotting. Cleaved CASP-3 was not detected on SHAM animals and only detected in the infarct area of AMI groups, but no difference was observed between them (Fig 6).

**Fig 6.**
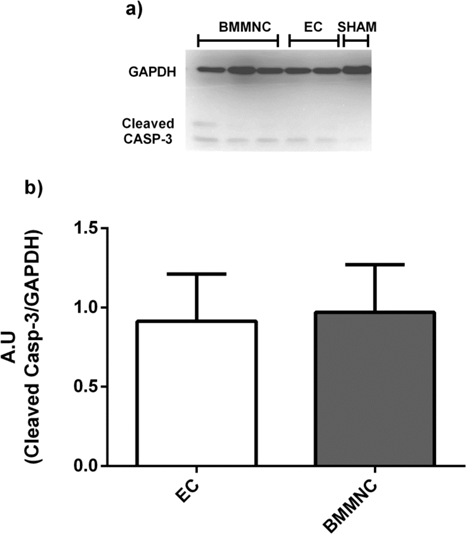
Cleaved caspase-3 levels. A) Western blot image showing the band for cleaved caspase-3 (lower band), normalized by GADPH (upper band). B) Comparison of cleaved caspase-3 levels in the EC and BMMNC groups. Levels were measured in arbitrary units (A.U.), which represent the band density of cleaved caspase-3 divided the by band density of the housekeeping protein GAPDH. Data expressed as mean ± SD. Statistical significance accepted at p<0.05.

## Discussion

Mononuclear cell-based therapy has emerged as one of the most optimistic potential treatments for cardiovascular diseases [12]. BMMNCs are composed of a diverse array of cells, but few are stem cells [29,30]; as a consequence, their therapeutic effect has been attributed to a poorly defined paracrine action. Here, we tested the hypothesis that this putative paracrine effect of BMMNCs could control the cardiac remodeling process after experimental AMI in rats.

In this study, we decided to encapsulate BMMNC in 1.5% sodium alginate and poly-L-Lysine, which prevents cells from interacting with the surrounding tissues but keeps them permeable to nutrients and metabolites. Capsules containing GFP^+^ BMMNCs were delivered all over the infarcted heart - with disrupted pericardium - shortly after coronary ligation and withdrawn after euthanasia (7 days after AMI).

In previous experiments we attempt to deliver BMMNC intra-myocardium but it cause the formation peri-capsular fibrosis (Supplementary Fig 4), thus we decided to delivered BMMNC capsules in the thoracic cavity based on the work of Lopes et al. (2014) that delivered encapsulated platelets in abdominal cavity for the treatment of liver failure [26]. At the end of the experiments, the capsules delivered intra-thoracic were still in contact with myocardium – with some spread out the thoracic cavity – and the majority of the cells were still viable, as evidenced by GFP^+^ BMMNCs visualized by fluorescence microscopy 7 days after AMI.

Nevertheless, this treatment was unable to prevent deleterious cardiac remodeling, as demonstrated by echocardiography findings in animals that received BMMNC compared with those that received empty capsules. In fact, an apparent worsening in ejection fraction was observed in animals in the BMMNC group, possibly due to a slightly, although not significantly, larger infarct size in animals allocated to this group. To confirm this finding, we decided to evaluate the progression of remodeling by comparing the variation of ejection fraction between days 2 and 7. This approach revealed no difference between groups, i.e., both groups had the same variation in ejection fraction over time. This change in LVEF is typical of the cardiac remodeling process.

It is important to highlight that the study protocol was designed with a 7-day follow-up period on the basis of previous data showing that major changes in the post-AMI heart occur within this period [31]. Additionally, all AMI groups began the protocol with the same infarct size and cTnI levels, as measured 48 h after AMI induction. Animals enrolled in the treatment groups had at least 30% infarct size with high cTnI levels (> 4.8 ng/mL, predictive value for AMI in rats) [32] and decreased LVEF.

As expected, 7 days after AMI, the morphometric parameters characteristic of cardiac remodeling were observed in the EC-treated group, namely: anterior wall thinning, increased left ventricular diameter, and decreased LVEF. No beneficial effect of BMMNC therapy on these parameters was observed in the cell-treated group.

Despite this lack of effect on the cardiac remodeling process, BMMNC therapy may still have modulated inflammatory cytokines. In fact, in many experimental studies showing beneficial outcomes of BMMNC therapy, these benefits were associated with modulation of inflammatory response, decreased oxidative stress, and increased vascularization secondary to release of cytokines and growth factors [33-36].

It is well known that IL-6 and TNF-α levels increase after AMI, a phenomenon associated with myofibroblast infiltration and tissue healing and fibrosis [37,38]. Unsurprisingly, the infarct area exhibited a threefold increase in IL-6 and TNF-α levels compared to the peri-infarct and remote areas. Again, no difference in these parameters was observed between animals treated with encapsulated BMMNCs and empty capsules.

BMMNCs have also been reported to produce anti-inflammatory cytokines, such as IL-1ra. Therefore, we analyzed levels of the anti-inflammatory cytokine IL-10. The main mechanism of action of this cytokine appears to involve control of the JAK-STAT pathway, leading to activation of the suppressor of cytokines signaling 3 (SOCS-3) and diminishing production and action of inflammatory cytokines such as IL-6 and TNF-α [39]. However, in our sample, there was no change in IL-10 levels, neither in the different regions of the heart nor between the treatment and control groups.

Finally, we analyzed activity of catalase, which is indicative of ability to counteract the production of reactive oxygen species (ROS) [40], and levels of cleaved CASP-3, a major marker of apoptotic signaling [41]. Again, treatment with encapsulated BMMNCs was not associated with any change in catalase activity or cleaved-CASP 3 levels, whether within the different areas of the heart or between the treatment and control groups.

Our results show a lack of any paracrine effect of BMMNCs 7 days in this rat model of experimental AMI. Nevertheless, ruling out the use of these cells as a regenerative treatment might be too hasty. Several factors can make it difficult to compare studies of BMMNC therapy for AMI, including timing of therapy, BMMNC extraction method, delivery sites, and preconditioning, which can influence both the quantity and quality of cells delivered for treatment. Such differences in studies make comparison of findings challenging, and, most importantly, hinder estimation of their potential translatability to clinical use.

Despite the controversial nature of this treatment, more than 30 clinical trials have been conducted [42,29], and some are still ongoing. Intriguingly, among all double-blind randomized clinical trials conducted to date, none has met primary endpoints, while studies that followed less rigorous protocols have found positive results [43]. Thus, protocol design and consistency of preclinical studies are key factors for determining the mechanism of action of BMMNCs and their applicability and translation to cardiovascular treatment.

Very few experimental animal studies employ randomization and blinded analysis, which may explain why many preclinical results are discredited in clinical trials. Our experimental protocol was carefully designed to follow the ARRIVE guideline [44], which advocates for randomization of animals into groups and blinded analysis performed by trained technicians. Unsurprisingly, our results seem very similar to those of previous randomized, double-blind clinical trials.

It is important to stress that our data must be interpreted in regard of the proposed hypothesis and considering the strengths and limitations of the experimental protocol. For instance, the permanent AMI model used herein is by far the most robust experimental model, but does not mimic human coronary artery disease, which is the leading cause of myocardial infarction worldwide. Another limitation is that BMMNC-loaded capsules were delivered loosely into the intrathoracic cavity; thus, paracrine secretion may have found different ways and fates. The use of bio-patches to keep capsules in contact with the infarct area could improve homing of paracrine secretion to cardiomyocytes, as previously described [45].

In short, we conclude that the putative paracrine action of encapsulated BMMNCs lacks efficacy to modulate events associated with post-infarct cardiac remodeling in a rat model of AMI.

## Acknowledgement

We like to give special thanks for Dr. Valeska Lagranha for her invaluable help and all the staff from UAMP and UEA at the Hospital de Clínicas de Porto Alegre for all support.

## Disclosure

The authors declare that they have no conflict of interest.

The protocol was carry-out in accordance to Guide for the Care and Use of Laboratory Animals^[46]^, the ARRIVE guideline^[47]^ and it was approved by the Ethics Committee on Animal Use from the Hospital de Clínicas de Porto Alegre (CEUA/HCPA) and registered with number 10-0246.

No human studies were carried out by authors for this article.

